# Biological activity of a stable 6-aryl-2-benzoyl-pyridine colchicine-binding site inhibitor, 60c, in metastatic, triple-negative breast cancer

**DOI:** 10.1101/2024.05.22.595349

**Authors:** Damilola Oluwalana, Kelli L Adeleye, Raisa I Krutilina, Hao Chen, Hilaire Playa, Shanshan Deng, Deanna N Parke, John Abernathy, Leona Middleton, Alexandra Cullom, Bhargavi Thalluri, Dejian Ma, Bernd Meibohm, Duane D Miller, Tiffany N Seagroves, Wei Li

## Abstract

**Background:** Improving survival for patients diagnosed with metastatic disease and overcoming chemoresistance remain significant clinical challenges in treating breast cancer. Triple-negative breast cancer (TNBC) is an aggressive subtype characterized by a lack of therapeutically targetable receptors (ER/PR/HER2). TNBC therapy includes a combination of cytotoxic chemotherapies, including microtubule-targeting agents (MTAs) like paclitaxel (taxane class) or eribulin (vinca class); however, there are currently no FDA-approved MTAs that bind to the colchicine-binding site. Approximately 70% of patients who initially respond to paclitaxel will develop taxane resistance (TxR). We previously reported that an orally bioavailable colchicine-binding site inhibitor (CBSI), VERU-111, inhibits TNBC tumor growth and treats pre-established metastatic disease. To further improve the potency and metabolic stability of VERU-111, we created next-generation derivatives of its scaffold, including 60c.

**Results:** 60c shows improved in vitro potency compared to VERU-111 for taxane-sensitive and TxR TNBC models, and suppress TxR primary tumor growth without gross toxicity. 60c also suppressed the expansion of axillary lymph node metastases existing prior to treatment. Comparative analysis of excised organs for metastasis between 60c and VERU-111 suggested that 60c has unique anti-metastatic tropism. 60c completely suppressed metastases to the spleen and was more potent to reduce metastatic burden in the leg bones and kidney. In contrast, VERU-111 preferentially inhibited liver metastases and lung metastasis repression was similar. Together, these results position 60c as an additional promising CBSI for TNBC therapy, particularly for patients with TxR disease.

## 1. Introduction

Breast cancer is the second-leading cause of cancer-related deaths in women, with >43,000 deaths expected in the US in 2023 [1]. Breast cancers are classified into four major subtypes based on expression of hormonal receptors (estrogen receptor, ER) or progesterone receptor, PR), and human epidermal growth factor 2 (HER2) [2, 3]. Hormone-receptor positive (HR+) and HER2+ breast cancers are primarily treated with endocrine therapy, CDK 4/6 inhibitors, or anti-HER2 antibodies conjugated to chemotherapy warheads, known as antibody-drug conjugates (ADCs) [4]. Triple-negative breast cancers (TNBC) remain the most challenging to treat. Only one targeted therapy is approved for metastatic TNBC, a TROP2-antibody conjugated to sactizumab govitecan [5, 6]. First-line TNBC treatment relies on a combination of cytotoxic chemotherapy drugs, including anthracyclines, cyclophosphamides, and anti-tubulin agents [6].

Tubulin inhibitors disrupt the dynamics of microtubule networks, causing mitotic arrest and triggering apoptosis [7]. The tubulin macrostructure is composed of two subunits, alpha- and beta-tubulin, which dynamically disassemble and reassemble as cells progress through the cell cycle. Disrupting tubulin networks generally leads to a G2/M block [7, 8]. There are at least seven identified microtubule-binding sites, including the colchicine, vinca and taxane sites [9, 10]. Paclitaxel (Taxol) binds to the taxane-binding site on beta-tubulin, stabilizing tubulin networks [11].

Although initially effective for TNBC, taxane use is associated with dose-limiting toxicity, poor aqueous solubility, peripheral neuropathy, and rapid onset of chemoresistance in some patients [12, 13]. Overexpression of the ATP-dependent efflux pumps, including P-glycoprotein (P-gp) or the β-tubulin III isoform [14, 15] is a primary mechanism of taxane resistance. Conversely, the colchicine binding site inhibitors (CBSIs), which prevent tubulin polymerization, destabilizing tubulin networks, can bypass taxane resistance [16–18]. We previously reported on the potency and efficacy of several unique CBSIs in various solid cancer types [19–27], including metastatic TNBCs [16, 28].

CBSIs based upon the 2-aryl-4-benzoyl-imidazole (ABI) scaffold are potent anticancer agents [29]. ABI-231, or VERU-111, is an orally bioavailable CBSI that was well-tolerated in phase I/II studies [30]. VERU-111 has anti-tumor and anti-metastatic activity in multiple pre-clinical models [16, 31]. New ABI class derivatives were made to improve the potency and metabolic stability of VERU-111, including CH-2-77, a 6-aryl-2-benzoyl-pyridine (ABP) with improved potency *in vitro* over VERU-111 [20]. CH-2-77 inhibits growth of treatment-naïve TNBC and taxane-resistant melanoma [20, 28]. However, CH-2-77 is metabolically labile, with a half-life of ∼10.8 minutes in human liver microsomes, limiting its use clinically [21]. Structure-activity relationship analysis was used to generate new CH-2-77 analogs that retained ABP scaffold potency while improving metabolic stability, resulting in the generation of compound 60c [21]. Like CH-2-77, 60c was effective in a taxane-resistant melanoma model [20]. We have shown that CH-2-77 represses metastatic breast tumor progression in taxane-sensitive models [28]; however, CH-2-77 efficacy in TxR breast cancer was not previously tested.

Herein, we compared the efficacy of 60c to CH-2-77 across multiple taxane-sensitive and TxR TNBC models, and performed a head-to-head comparison of 60c versus paclitaxel in the highly metastatic, taxane-refractory HCI-10-Luc2 TNBC PDX model [16, 32, 33]. In vitro, 60c is as potent as CH-2-77 in taxane-sensitive and TxR TNBC cells, with increased potency relative to VERU-111. 60c represses primary tumor growth in MDA-MB-231-TxR xenografts, and in the HCI-10-Luc2 PDX. Because the HCI-10-Luc2 PDX generates widespread metastases to multiple organs within the same animal, we could observe organ tropism for metastatic suppression comparing 60c and VERU-111. Whereas 60c more efficiently repressed spleen, kidney, and bone metastasis than VERU-111, VERU-111 inhibited liver metastasis more efficiently. Overall, these results demonstrate that 60c is an effective CBSI drug candidate for either taxane-sensitive or taxane-refractory TNBC patients.

## 2. Materials and Methods

### 2.1 Chemicals, cell culture, and PDX materials

Paclitaxel, colchicine and combretastatin A4 phosphate, CA-4P (also known as fosbretabulin) were purchased from LC Laboratories (P-9600, purity 99.5%, Woburn, MA, USA), Sigma-Aldrich (C9754, purity >95.0%, St. Louis, MO, USA) and MedKoo Biosciences (200800, purity >98%, Morrisville, NC), respectively. 60c, VERU-111, and CH-2-77 were synthesized as in [20, 21, 26]. MDA-MB-231 (231), MDA-MB-468 (468) and Hs578T TNBC cell lines were obtained from the American Tissue Culture Collection (ATCC) and SUM159 cells were obtained from Dr. Bisrat Debeb at MD Anderson Cancer Center. TNBC-TxR sublines were generated by gradually increasing the concentration of paclitaxel on surviving cells until a drug resistance index (RI) of ≥ 10 was achieved, as in [16, 28]. Parental and TxR TNBC cell lines were grown in DMEM-Hi (231), DMEM/F12 (468, SUM159) or RPMI (Hs578T) supplemented with 10% FBS and 1% antibiotic-antimycotic solution. SUM159 cells were supplemented with 1 μg/mL of insulin and hydrocortisone, and Hs578T cells\ with 10 μg/mL of insulin. All cells were authenticated at the University of Arizona Genetics Core (RRID:SCR_012429) before beginning experiments, and were screened for mycoplasma (MycoAlert kit, Lonza, Basel, Switzerland). HCI-10 PDX tumor fragments were provided by Dr. Alana Welm and the Huntsman Cancer Institute (HCI) Pre-Clinical Research Shared Resource [33]. Generation of the HCI-10-Luc2 luciferase-labeled TNBC PDX subline was previously described in [34], and taxane-resistance shown in [16]. HCI-10-Luc2 PDX-derived cells were cultured in M87 growth media containing 2% FBS [32].

### 2.2 Acetylated α-tubulin assay

231 cells grown to 80% confluence were incubated with increasing concentrations of 60c (4, 8, and 16 nM) or 16 nM of VERU-111 or CH-2-77 for 24 h. Whole cell extract (20 μg) was resolved on 4-15% gradient gels, transferred to PVDF membrane, incubated overnight at 4°C with anti-α-tubulin antibodies (Cell Signaling Technology, CST, #12152), followed by incubation with IgG-HRP-conjugated antibodies (CST #7076) at RT for 1 h, and detection by SuperSignal West Pico PLUS substrate (Thermo-Scientific). GAPDH (rabbit anti-GAPDH-HRP, CST #3683) was used as a loading control. Blots were imaged by the Bio-Rad ChemiDoc MP system.

### 2.3 EBI tubulin crosslinking assay

231 cells grown to 80% confluence were incubated with increasing concentrations of VERU-111, 60c, or CH-2-77 (100 nM, 300 nM, 1 μM) or 1 μM colchicine for 2 h. EBI [N,N’-ethylene-bis(iodoacetamide),100 μM] was added and incubated for 1.5 h. Protein was resolved and detected as in 2.2. Membranes were incubated overnight at 4°C with anti-β-tubulin (CST #2128), followed by IgG secondary HRP-conjugated antibody (CST #7074). GAPDH was used as a loading control.

### 2.4 Cell confluence and IC_50_ growth inhibition assays

Cells were seeded overnight into 96-well plates (3,000 cells/well; n = 8), treated with increasing concentrations of 60c, CH-2-77, VERU-111 or paclitaxel, and imaged with the IncuCyte S3 live-cell imaging system (Sartorius, Göttingen, Germany). Phase-masking algorithms were applied to determine cell confluence. Cells were plated similarly for MTS assays, an indirect measure of growth inhibition, but over a wider concentration range. Drug response was first normalized to untreated vehicle controls and/or to initial seeding density, data were converted to a log scale, then plotted using non-linear regression best fit analysis in GraphPad Prism 9.0. The mean IC_50_ ± SEM was derived from at least 3 biological replicates.

### 2.5 Colony formation assay

Cells were plated into 96-well plates at low density (500 cells/well; n = 8) and treated 24 h later with increasing drug concentrations. Media was replaced every 5 days, and images obtained every 12 h using the IncuCyte S3. The phase-masking algorithm was applied to enumerate colony formation. Data were exported to Prism 9.0 and normalized to seeding density. Cells were washed with PBS, fixed with methanol, stained with 0.5% crystal violet, and imaged at endpoint.

### 2.6 Wound healing assay

Wound healing assays were performed as in [26, 35]. Post-wounding, cells were treated with increasing concentrations of each compound until wound closure was observed for the vehicle control. Wound width was quantitated using IncuCyte software. Data are representative of two biological replicates (n = 4 replicates/treatment/cell line).

### 2.7 Protein extraction and western blotting

Cells grown to 80% confluence were incubated with increasing concentrations of 60c (4, 8, and 16 nM) or 16 nM of VERU-111 or CH-2-77 for 24 h. Whole cell lysates (35 μg) were resolved on 4-12% Bis-Tris gels and transferred onto PVDF. Blots were exposed to primary antibodies against cleaved-PARP, phosphorylated BCL2, or cleaved caspase-3 (CST #5625, #2827, #9661) at 4°C overnight. After washing, membranes were incubated with anti-rabbit whole IgG secondary HRP-conjugated secondary antibodies (Jackson ImmunoResearch, West Grove, PA) at RT for 1 h and detected by ECL reagent (WBKLS0500, Millipore), then exposed to film.

### 2.8 Transwell migration and invasion assays

231-TxR cells were grown to 80% confluence, then serum-starved for 24 h. Cells (20,000) were plated into the upper chamber of transwell migration chambers (Corning #353097) or matrigel-coated chambers (Corning #354480). Cells were incubated with paclitaxel, 60c, or CH-2-77, and chemoattracted to growth medium containing 10% FBS. HCI-10 PDX-derived cells were grown to 80% confluence, plated at 80,000 cells/chamber and chemoattracted to growth medium containing 50 nM of human HGF (Life Technologies). Drugs were added in transwell assays at the time of cell seeding. After 24 h, membranes were fixed, stained, imaged and quantified using ImageJ software.

### 2.9 Orthotopic MDA-MB-231-TxR xenografts

All animal studies adhered to NIH Principles of Laboratory Animal care and protocols approved by the local Institutional Animal care and Use Committee at UTHSC (protocols 20-0397 or 20-0181). Female NSG mice (5 - 6 weeks old, Jax Labs, stock#005557) were inoculated with 2.5 × 10^5^ 231-TxR cells suspended in 10 µL HBSS in each inguinal mammary fat pad (m.f.p.) using a Hamilton syringe with a 1/2’’ 26G PT2 needle. Tumors were measured with digital calipers and volume was calculated by (width^2^ x length)/2. Once one tumor per mouse grew to 100 mm^3^, mice were randomized such that mean tumor volumes were equivalent across treatment cohorts [**1**: vehicle (ethanol:Cremophor-EL:0.9%saline = 1:1:18, 3x/week, IP, n = 6 mice); **2**: 10 mg/kg paclitaxel (3x/ week, IP, n = 6 mice); **3**: 35 mg/kg CA-4P (0.9% saline, 5x/week, IP, n = 7 mice); **4**: 25 mg/kg CH-2-77 (PEG300:Tween80:water = 4:1:10 ratio; 5x/week, IP, n = 6 mice); **5**: 25 mg/kg 60c (PEG300:Tween80:water = 3:1:6 ratio; 5x/week, IP, n = 7 mice)]. Since CA-4P is a prodrug, a 35 mg/kg dose yields 25 mg/kg active combrestatin *in vivo* for equivalent dosing with other CBSIs. Any tumors not ∼100 mm^3^ at the time of cohort assignment were excluded from analysis. Excised tumors were photographed at endpoint, and tumor volumes re-calculated by caliper measurement. Tumors were bisected and one half fixed overnight in 10% neutral-buffered formalin (NBF) before paraffin embedding and sectioning. All other viable tumor tissue was collected and flash-frozen in liquid nitrogen. Lungs were harvested after inflation with saline, and then fixed overnight in 10% NBF for histopathology.

### 2.10 HCI-10-Luc2 PDX model and bioluminescent imaging

PDX tumor fragments (∼2 mm^3^) harvested from a tumor-bearing “donor” mouse were bilaterally implanted into each inguinal m.f.p. of recipient NSG females (6 - 7 weeks old). Treatment cohorts were: [**1**: vehicle (ethanol:Cremophor-EL:0.9%saline = 1:1:18, 3x/week, IP, n = 9 mice); **2**: 10 mg/kg paclitaxel (3x/week, IP, n = 9 mice); **3**: 20 mg/kg 60c (PEG300:Tween80:water = 3:1:6 ratio; 5x/ week, IP, n = 9 mice)]. Mice were randomized before assignment to a treatment arm as in 2.9, such that median tumor volume (∼80 mm^3^) and bio-imaging photon flux were equivalent among cohorts. Animals were euthanized and organs processed as in [16]. For bioluminescent imaging, mice were injected with D-luciferin salt 5 minutes before anesthesia, then imaged in the IVIS Lumina XRMS instrument 10 minutes post-injection. At study endpoint, each intact mouse was imaged, then tissues rapidly harvested and individually imaged *ex vivo*. Signal counts were converted to total photon flux using LivingImage software as in [16].

### 2.11 Histology and immunohistochemistry (IHC) analysis

Paraffin-embedded specimens were prepared as previously described [31]. Slides were incubated with primary antibodies to either phospho-histone H3(Ser10), cleaved caspase-3, or CD31 (1:200, CST, #9701, #9661, or #77699) overnight in humid chambers at 4°C. Slides were developed as described in [16] and imaged with a Pannoramic 250 slide scanner (3DHISTECH, Budapest, Hungary). Lung metastasis was evaluated after IHC with an anti-human-specific mitochondrial antibody (1:1,000, Abcam, #ab92824) Automated quantification of phospho-H3, CD31, and cleaved caspase-3 was performed as in [16] and the H-scores are reported.

### 2.12 Drug distribution analysis

Female NSG mice aged 10-12 weeks were dosed with 25 mg/kg 60c intraperitoneally, or 25 mg/kg VERU-111 orally. Mice were sacrificed after 30 minutes (60c) or 1 hour (VERU-111), respectively, time based on [21, 36]. Blood was collected into heparin-coated tubes (BD#365985) and placed on ice, and whole-body perfusion was performed with 0.9% saline. Plasma was collected and frozen. All tissue samples were washed twice in PBS, dried, and weighed, then suspended in 4 mL of PBS/mg tissue weight for homogenization. Homogenate (50 μL) was diluted into 200 μL of ice-cold methanol and samples centrifuged for 10 minutes at 12,000 rpm. The supernatant was then collected for analysis by LC-MS/MS using procedures as in [21].

### 2.13 Statistical analysis

Unless otherwise described, all raw data were entered into GraphPad Prism 9.0 and analyzed using one-way or two-way ANOVA followed by multiple pairwise comparison tests (Student’s *t*-tests), and the mean ± SEM is shown. Significance was determined using a 95% confidence threshold.*p*-values are indicated by asterisks defined as, \**p*<0.05, \*\**p*<0.01, *** *p*<0.001, **** *p*<0.0001.

## 3. Results

### 3.1 60c binds more efficiently to tubulin than VERU-111 or CH-2-77

Analogs of VERU-111 were made to improve potency, including CH-2-77. Analogs of CH-2-77 were then made to improve metabolic stability. Specifically, the 60c analog was derived from CH-2-77 by incorporating a fluorine in the “B”-ring of the CH-2-77 scaffold (**Fig. 1A**). Although 60c retained similar potency as CH-2-77, it had enhanced metabolic stability relative to CH-2-77 in human liver microsomes (*t*_1/2_= 29.4 ± 2.4 min vs. 10.8 ± 0.6 min, respectively) and mouse liver microsomes (*t*_1/2_= > 4 h vs. 60 ± 1.8 min, respectively) [21].

**Figure 1.**
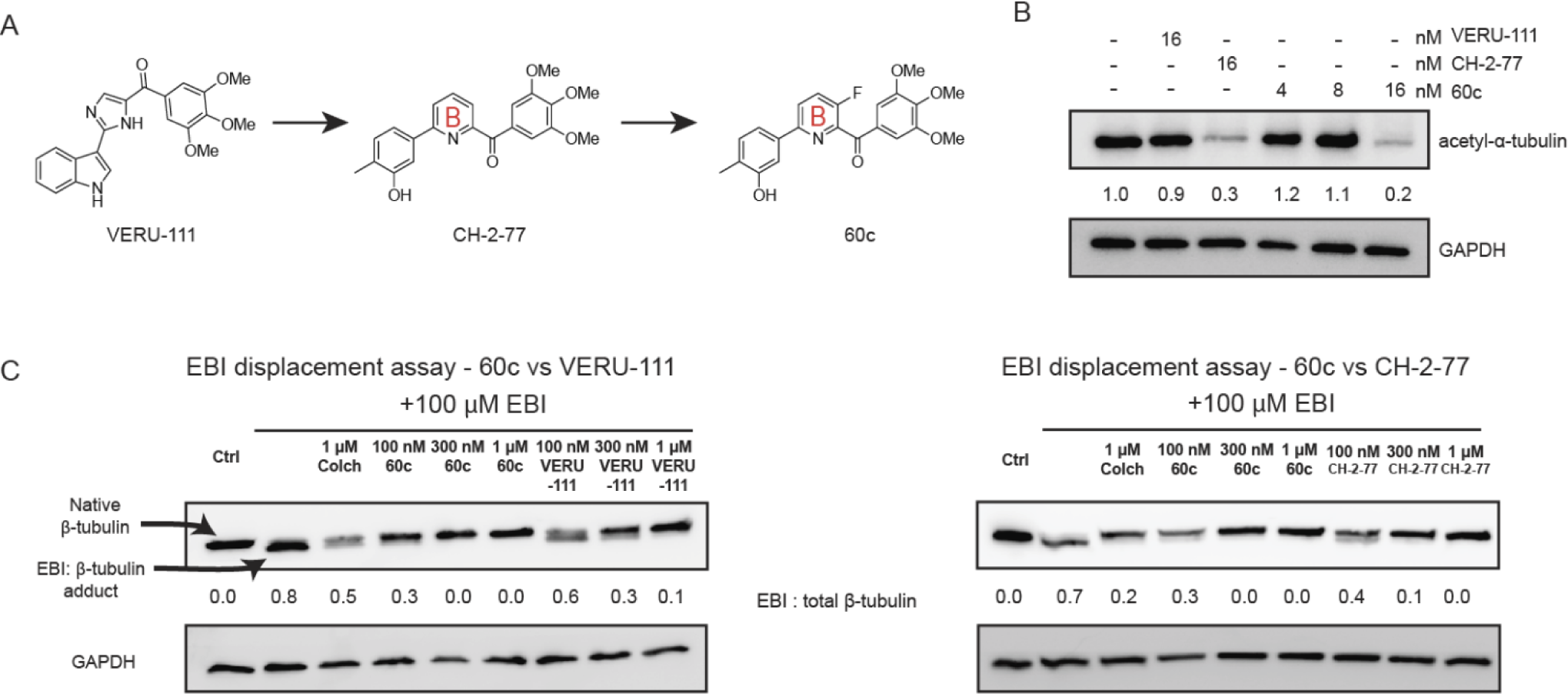
Evolution of next-generation VERU-111 analogs, which bind with higher efficiency to the colchicine-binding site (CBSI) of β-tubulin. (**A**) Chemical structures of VERU-111 (the parent scaffold, formerly ABI-231), CH-2-77 (formerly known as 4v), and 60c, derived from CH-2-77. (**B**) Acetylated α-tubulin levels are expressed relative to the vehicle control. (**C**)The EBI competitive binding assay was used to compare the efficiency of tubulin binding by the CBSIs over a range of 100 nM to 1 mM) relative to colchicine (1 mM). The ratio of the EBI-adduct to total β-tubulin is indicated below each treatment.

Acetylation of α-tubulin promotes microtubule stability [37–39]. Acetylated α-tubulin signal is lost after treatment with 60c and CH-2-77 (16 nM), but signal for VERU-111 (16 nM) is comparable to the vehicle control, indicating more efficient tubulin destabilization with the next generation CBSIs (**Fig. 1B**).

To compare the β-tubulin binding affinity between VERU-111, 60c, and CH-2-77, the EBI assay was used. EBI crosslinks two cysteine residues of β-tubulin within the colchicine-binding site [40], causing EBI-adducts to migrate faster than native β-tubulin by western blotting. If compounds are more tightly bound to β-tubulin than EBI, more native β-tubulin protein is detected. At equivalent concentrations, 60c outcompetes EBI more efficiently than VERU-111 or CH-2-77 (**Fig. 1C**). The adduct completely disappears in the presence of 100 nM 60c but is still present after exposure to 300 nM of VERU-111 or 100 nM of CH-2-77. Overall, both 60c and CH-2-77 have enhanced β-tubulin binding and destabilize tubulin more efficiently than VERU-111.

### 3.2 60c is equipotent to CH-2-77 to inhibit cell growth and cell survival in taxane-sensitive or taxane-resistant TNBC cells

We assessed the anti-proliferative effect of 60c relative to VERU-111 and CH-2-77 in a panel of four TNBC models (231, Hs578T, SUM159, and 468) and their TxR sublines. In both parental and TxR lines, 60c inhibits growth in the low nanomolar range (4 – 10 nM) with comparable potency to CH-2-77 as assessed by an MTS assay (**Table 1 and Supp. Fig. 1**). In a microscopy-based growth assay measuring cell confluence, both analogs were more potent than an equivalent concentration of VERU-111 in 231-TxR, Hs578T-TxR and SUM159-TxR cells, and were equipotent to VERU-111 in 468-TxR cells (**Fig. 2A-D and Supp. Fig. 2**). As expected, each TxR TNBC subline resisted paclitaxel treatment (**Table 1**). 60c was comparable to CH-2-77 to inhibit colony growth at study endpoint (**Supp. Fig. 3**) or over time (**Supp. Fig. 4**) in a dose-dependent manner. In contrast, paclitaxel did not inhibit clonogenic growth in the TxR cells.

**Figure 2.**
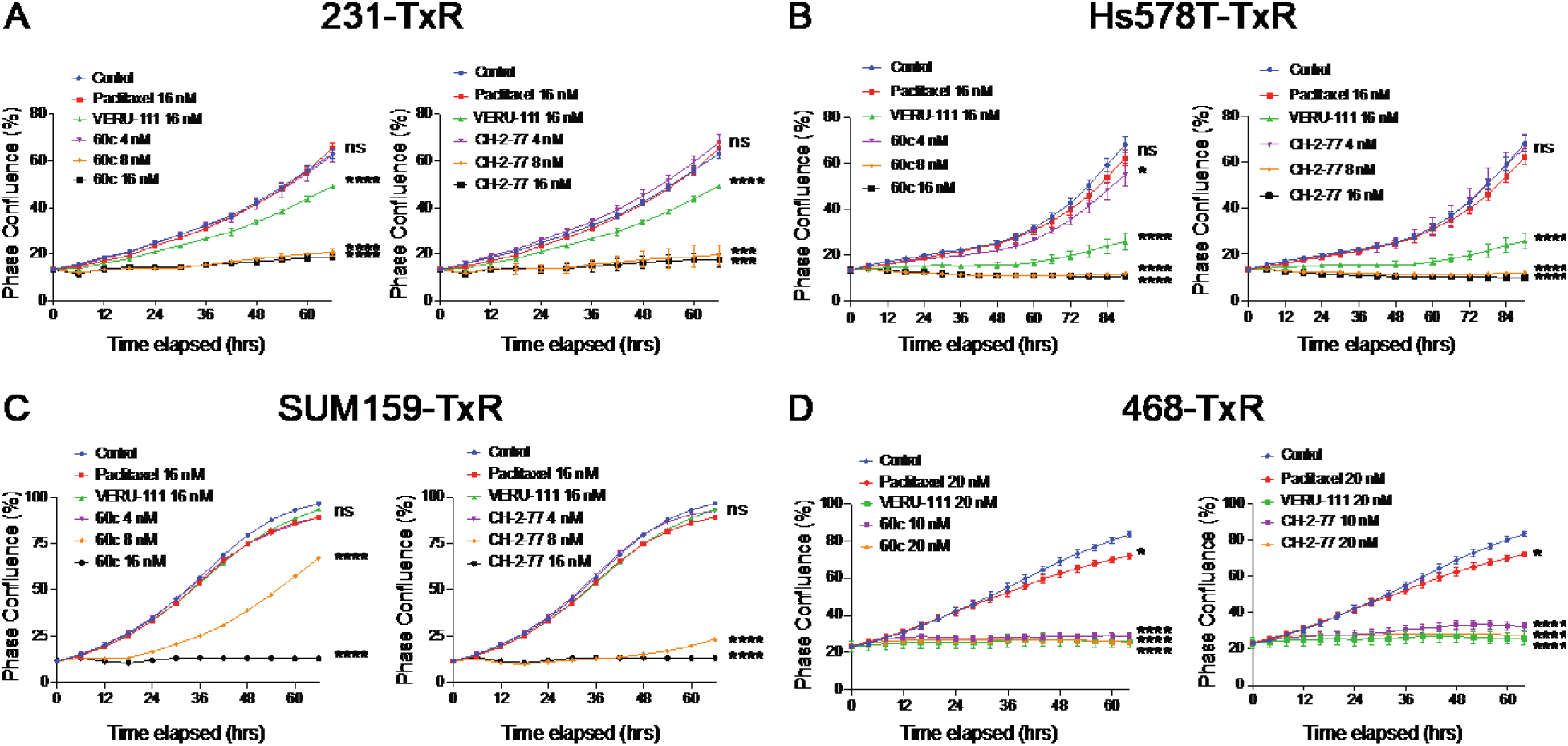
Dose-dependent inhibition of breast cancer cell growth by CH-2-77 or 60c versus paclitaxel in a panel of taxane-resistant TNBC cell lines, measured in real-time by cell confluence. Taxane-resistant (TxR) sublines of MDA-MB-231 (**A**), Hs578T (**B**), SUM159 (**C**) and MDA-MB-468 (**D**) cells were exposed to paclitaxel, VERU-111, 60c or CH-2-77 for 66 h (231, SUM159, and 468), or 90 h (Hs578T) over the course of each model’s log-phase growth period. Drugs were added at the indicated final concentration, and the medium was not changed over the course of the experiment. Cell growth was measured in real-time using the IncuCyte S3 imager and quantified using the percent (%) confluence algorithm. Representative drug response curves from at least three independent biological replicates/cell line are shown; *** *p* <0.001; **** *p* <0.0001; ns, not significant.

**Table 1.** IC_50_ (nM) ± SEM values for paclitaxel, VERU-111, 60c and CH-2-77 in a panel of taxane-sensitive and taxane-resistant TNBC lines, calculated at growth endpoints by the MTS assay. The grand mean of three independent biological replicates is shown. MDA-MB-231-TxR, Hs578T-TxR, SUM159-TxR and MDA-MB-468-TxR cells were selected to be resistant to 90 nM, 95 nM, 100 nM or 120 nM of paclitaxel, respectively.

| Cell Line | Paclitaxel (nM) | VERU-111 (nM) | 60c (nM) | CH-2-77 (nM) |
| --- | --- | --- | --- | --- |
| 231 | 4.3 ± 0.4 | 9.0 ± 2.6 | 6.0 ± 0.8 | 6.3 ± 0.5 |
| MB231-TxR | 140 ± 18 | 9.9 ± 1.2 | 4.6 ± 0.3 | 4.5 ± 0.4 |
| Hs578T | 4.1 ± 1.6 | 10.4 ± 3.1 | 2.8 ± 1.0 | 3.1 ± 0.1 |
| Hs578T-TxR | 56 ± 11 | 9.2 ± 1.1 | 3.5 ± 1.0 | 4.7 ± 2.2 |
| SUM159 | 6.2 ± 1.5 | 22.5 ± 3.1 | 4.3 ± 1.4 | 3.7 ± 0.1 |
| SUM159-TxR | 109.0 ± 4.4 | 22.1 ± 3.2 | 2.7 ± 0.1 | 3.0 ± 0.1 |
| 468 | 5.2 ± 0.2 | 15.6 ± 0.3 | 3.9 ± 0.8 | 5.0 ± 1.3 |
| 468-TxR | 354 ± 92 | 12.5 ± 4.4 | 3.1 ± 1.8 | 4.0 ± 2.0 |

### 3.3 CH-2-77 and 60c inhibit wound closure in taxane-resistant TNBC cells

231-TxR and Hs578T-TxR cells were treated with increasing concentrations of 60c and CH-2-77. Paclitaxel and VERU-111 (16 nM) were included as controls. In 231-TxR cells, both 60c and CH-2-77 suppressed wound closure at 8 nM, although 8 nM of 60c inhibited migration more effectively than 8 nM of CH-2-77 **(Fig. 3A)**. In Hs578T-TxR cells, 8 nM of CH-2-77 and 60c inhibited wound healing with similar efficiency (**Fig. 3B**). In contrast, in parental cells, 12 nM of CH-2-77 or 60c repressed wound closure with similar or greater efficacy than 16 nM paclitaxel, whereas treatment with 16 nM of VERU-111 had no significant effect on wound closure (**Supp. Fig. 5**).

**Figure 3.**
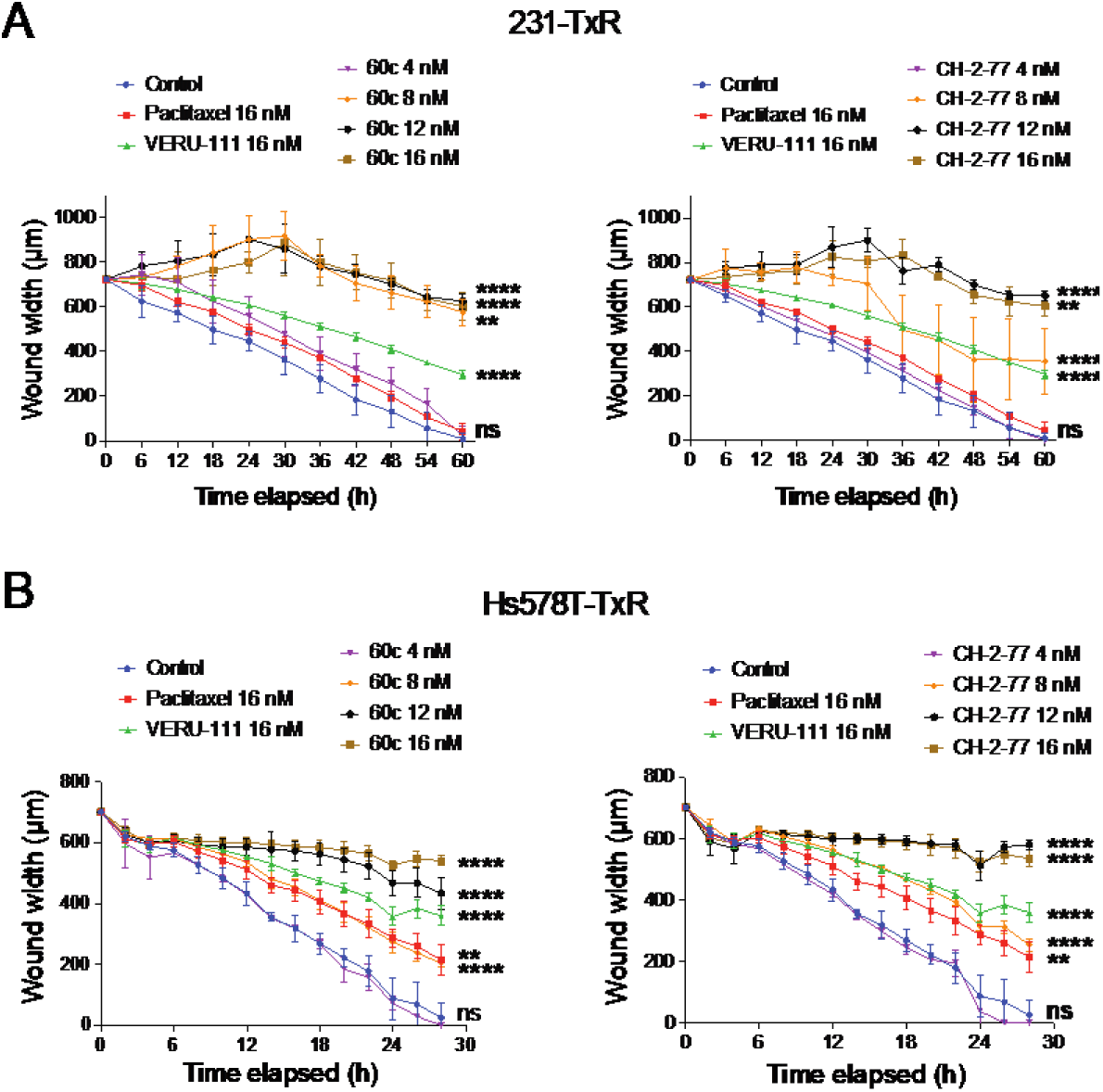
Dose-dependent inhibition of wound healing by 60c and CH-2-77 in taxane-resistant TNBC cells. Changes in wound width (μM) were measured in real-time using the IncuCyteS3 imager in wounded 231-TxR (**A**) and Hs578T-TxR (**B**) cells treated with paclitaxel (16 nM), VERU-111 (16 nM), or increasing concentrations of 60c and CH-2-77 (4, 8, 12 or 16 nM). Data were analyzed by two-way ANOVA followed by Dunnett’s multiple comparisons testing. All *p*-values are shown relative to the vehicle control at the experimental endpoint, when vehicle cells had filled the wound; ** *p* <0.01; *** *p* <0.001; **** *p* <0.0001; ns, not significant.

### 3.4 60c and CH-2-77 induce apoptosis and repress cell migration and invasion more potently than paclitaxel or VERU-111

We next assessed whether 60c induces apoptosis in 231 and Hs578T parental cells and TxR sublines. We previously showed that VERU-111 induces apoptosis in breast cancer cells [16, 31]. We screened for dose-dependent changes in cleaved-PARP, phosphorylated-BCL2, and cleaved caspase-3 by western blotting. 60c triggered apoptosis at a concentration of 8 nM in 231 parental and 231-TxR cells, and at 16 nM in Hs578T parental and TxR cells (**Fig. 4A**). Next, we investigated whether 60c reduces 231-TxR cell migration or invasion in a transwell assay. Both 60c and CH-2-77 significantly suppressed migration at 10 nM, whereas VERU-111 n(15 nM), and paclitaxel (10 nM) had no significant effect on migration (**Fig. 4B**). 60c and CH-2-77 suppressed invasion by ∼ 75% at a concentration of 10 nM. Of note, despite failing to repress cell migration, paclitaxel (10 nM) reduced the invasion of 231-TxR cells through Matrigel by ∼25%.

**Figure 4.**
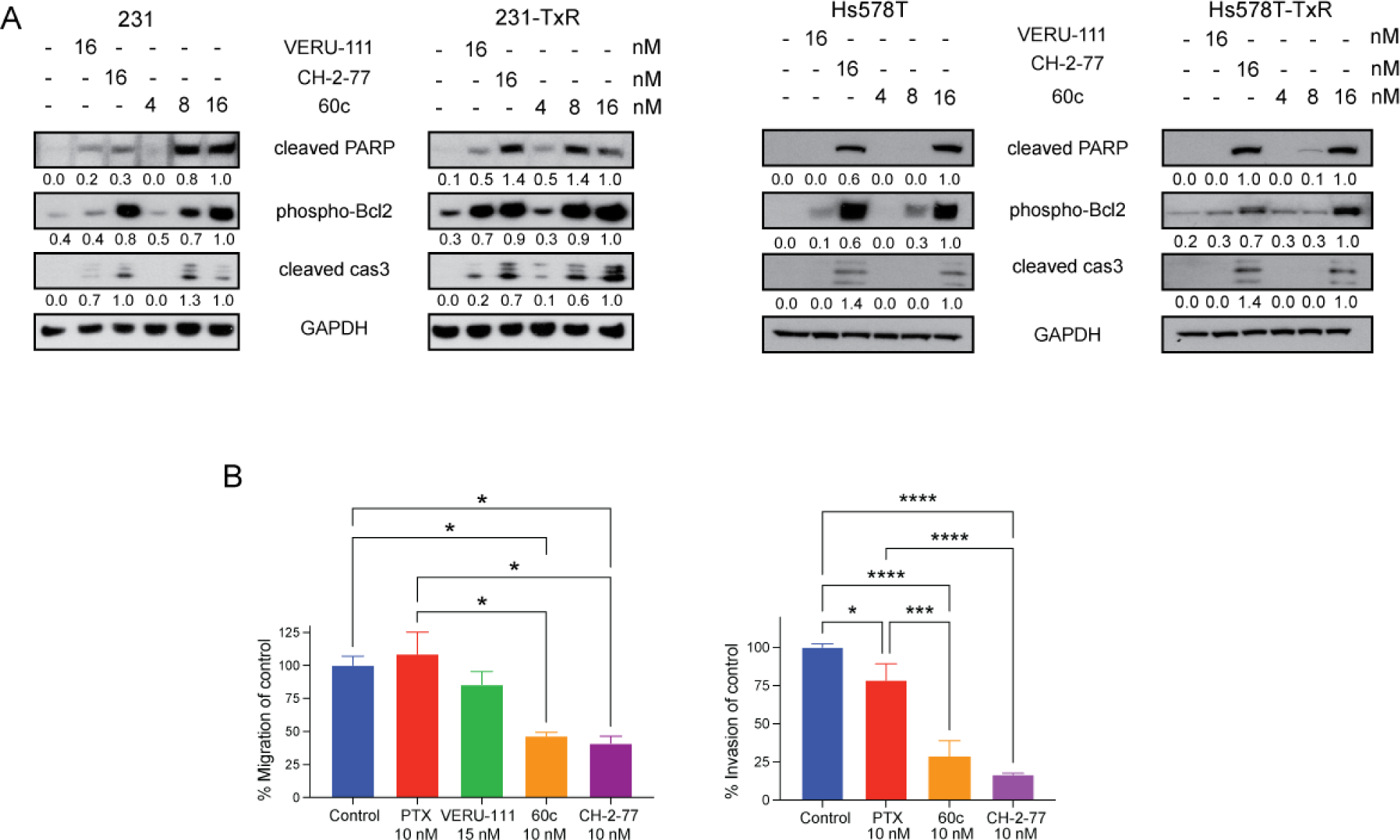
Comparison of CH-2-77 and 60c to induce apoptosis and to inhibit migration and invasion. (**A**) Changes in expression levels of apoptotic markers were compared by western blotting after 24h of exposure to VERU-111 (16 nM), CH-2-77 (16 nM) or increasing concentrations of 60c (4, 8, and 16 nM) in parental and TxR 231 and Hs578T cells. Expression levels of cleaved-PARP (RRID: AB_10699459), phosphorylated BCL2 (RRID: AB_659950), or cleaved caspase-3 (RRID: AB_2341188) were first normalized to Ponceau S staining and the 16 nM 60c-treated lane was set to 1.0. GAPDH (RRID: AB_1642205) serves as a loading control (**B**) The migration and invasion indices of 231-TxR cells (normalized to control) were determined in transwell assays after exposure of 60c or CH-2-77 (10 nM) for 24 h. Paclitaxel (10 nM) and VERU-111 (15 nM) were included as reference drugs controls for migration, and paclitaxel (10 nM) was used for invasion. Migration data are representative of two biological replicates (n = 3 technical replicates/assay) and invasion data were generated from three technical replicates in one assay. Data were analyzed by two-way ANOVA followed by Dunnett’s multiple comparisons testing. All *p*-values from pairwise comparisons are shown; * *p* <0.05; *** *p* <0.001; **** *p* < 0.0001.

### 3.5 60c, CH-2-77 and CA-4P are equipotent to suppress 231-TxR xenograft growth and lung metastasis

CH-2-77 inhibits growth of taxane-sensitive 231 xenografts and lung metastasis in a dose-dependent manner [28]. CA-4P, a CBSI in development for solid tumors, was also potent to inhibit metastatic breast cancer [28]. Since CH-2-77 bypasses taxane resistance [21], we used the 231-TxR xenograft model to compare the efficacy of CH-2-77 directly to 60c (each drug was administered 25 mg/kg, 5x/week IP), and to CA-4P (35 mg/kg, 5x/week, IP) and the standard of care chemotherapy, paclitaxel (10 mg/kg, 3x/week, IP).

As expected, 231-TxR tumors were refractory to a dose of paclitaxel that is highly effective against taxane-sensitive MDA-MB-231 xenografts [28]. Despite the improved metabolic stability observed for 60c over CH-2-77, we observed that all three CBSIs were equipotent to repress 231-TxR xenograft growth (**Fig. 5A**). Equally important, none of the compounds tested caused gross toxicity, including body weight loss (>10% of body weight, indicated by dashed lines in **Fig. 5B**). At study endpoint, no significant differences in mean tumor volume or tumor wet weight were observed among the CA-4P, CH-2-77 or 60c cohorts, whereas paclitaxel-treated tumors were equivalent in size and weight to the vehicle control treatment (**Fig. 5C-D**).

**Figure 5.**
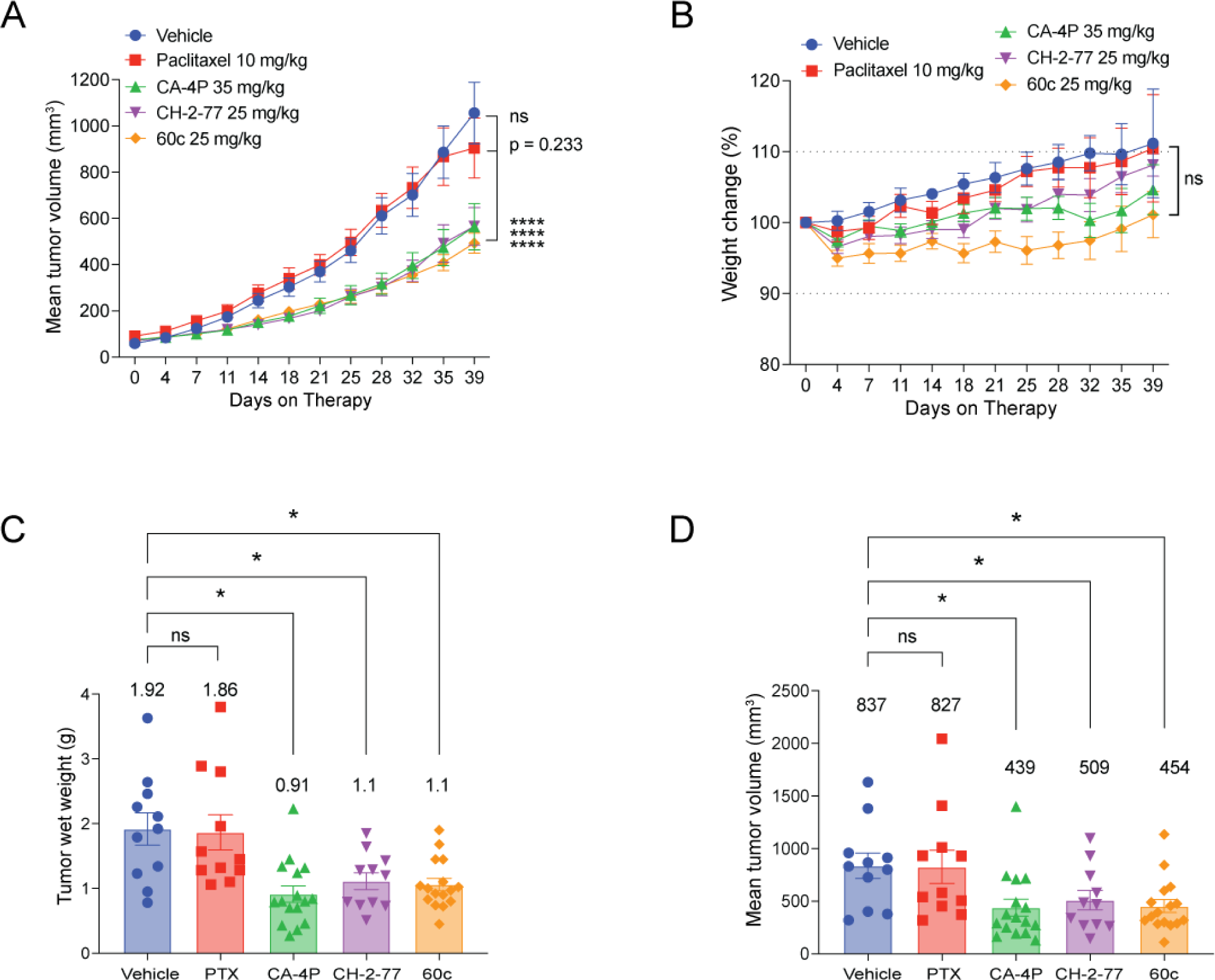
60c, CH-2-77 and CA-4P are equipotent to inhibit growth of taxane-resistant MDA-MB-231 tumors. (**A-B**) Mice bearing 231-TxR tumors were treated with either vehicle, paclitaxel (10 mg/kg, 3x/week, IP), CA-4P (35 mg/kg, 5x/week, IP), CH-2-77 (25 mg/kg, 5x/week, IP) or 60c (25 mg/kg, 5x/week, IP), head-to-head using equivalent doses by molecular weight (after correction for the processing of the CA-4P prodrug in vivo). Mean tumor volume (**A**) and the percent change in body weight (**B**) over the course of treatment are shown (± SEM). Tumor wet weight (**C**) and final tumor volume (**D**) of excised tumors was measured at study endpoint; individual tumors are plotted in a scatter plot wherein the bar indicates the mean and the error bars indicate SEM. The numerical mean is indicated above each cohort. All data were analyzed by one-way ANOVA and post-hoc Student’s *t*-tests with Welch’s correction; * *p* <0.05; **** *p* <0.0001; ns, not significant.

Because CBSIs induce cell death, H&E-staining was also used to compare necrosis of whole tumor sections between groups. Tumor necrosis was significantly increased among all treatment groups compared to vehicle control (**Supp. Fig. 6A**). Immunohistochemistry was performed to compare cell proliferation (phospho-histone-H3, Ser10), apoptosis (cleaved caspase-3) and microvessel density (CD31). Cell proliferation was potently reduced in response to CBSI treatment, with no significant differences among CA-4P, CH-2-77 or 60c (**Supp. Fig. 6B**). Likewise, the CA-4P and CH-2-77 -treated tumors showed more evidence of apoptosis than the paclitaxel group, while the data for 60c trended towards significance (*p*=0.09) (**Supp. Fig. 6C**).

Despite prior observations that CBSIs repress angiogenesis [17, 41, 42], no significant differences in microvessel density were observed among treatment groups (data not shown). Lung metastasis was evaluated as previously described [31]. CH-2-77, 60c and CA-4P each repressed lung metastasis with an ∼9-fold reduction in the H-score relative to vehicle treatment (**Supp. Fig. 6D**). Surprisingly, there was not a significant difference in metastasis burden observed between the paclitaxel and the CBSI drug treatment groups, indicating that in the context of metastasis, but not primary tumor growth, 231-TxR cells partially respond to paclitaxel therapy.

### 3.6 60c is highly effective to repress tumor growth and metastatic burden in a taxane-refractory, highly aggressive patient-derived xenograft model

The HCI-10-Luc2 TNBC model is a luciferase-labeled subline of a PDX developed from a patient whose cancer was refractory to a panel of conventional chemotherapies, including taxanes [33]. Luciferase labeling facilitates monitoring of primary tumor growth and metastatic spread simultaneously and longitudinally in the same animal. The HCI-10-Luc2 PDX efficiently metastasizes from the m.f.p. to axillary lymph nodes (AxLN), lungs, liver, bones, and kidney [16, 43]. We previously validated that the HCI-10-Luc2 is paclitaxel-refractory but is highly sensitive to VERU-111 [16].

We first determined the IC_50_ values of the next-generation CBSI analogs CH-2-77 and 60c in cells derived from the HCI-10-Luc2 PDX by the MTS assay, confirming that HCI-10-Luc2 cells are resistant to paclitaxel, but are highly sensitive to both CH-2-77 and 60c (**Table 2**; **Fig. 6A**). In transwell assays, CH-2-77 and 60c repressed PDX cell migration (**Fig. 6B**) and invasion (**Fig. 6C**). Since 60c has improved metabolic stability in liver microsomes over CH-2-77, we selected 60c to evaluate against paclitaxel in this model. Because of the longer duration of tumor growth of this PDX model relative to 231-TxR tumors, increasing the potential for cumulative toxicity, the 60c dosing regimen was revised downward to 20 mg/kg, 5x/week, IP, to achieve a cumulative dose of 100 mg/week vs. 125 mg/week in the 231-TxR model. Mice bearing 231-TxR tumors received ∼725 mg of 60c after 39 days of therapy, whereas mice bearing PDX tumors received ∼820 mg of 60c after 57 days of therapy. x 60c suppressed primary tumor growth by ∼3.5-fold, without body weight loss, while paclitaxel therapy was ineffective (**Fig. 6D-E**). By study endpoint, 60c reduced *ex vivo* tumor volume (**Fig. 6F**) and tumor wet weight (**Fig. 6G**) by ∼3.5-fold relative to the vehicle control. In contrast, there were no significant differences between the vehicle and paclitaxel-treated tumors during the study or at endpoint (**Fig. 6D**, **6F-G**), as expected. The reduction in primary tumor volume in the 60c cohort evident from images of excised tumors (**Fig. 6H**) was validated by bio-imaging, which revealed a 3.82-fold reduction in tumor photon flux between vehicle and 60c treatment (**Fig. 7A**).

**Figure 6.**
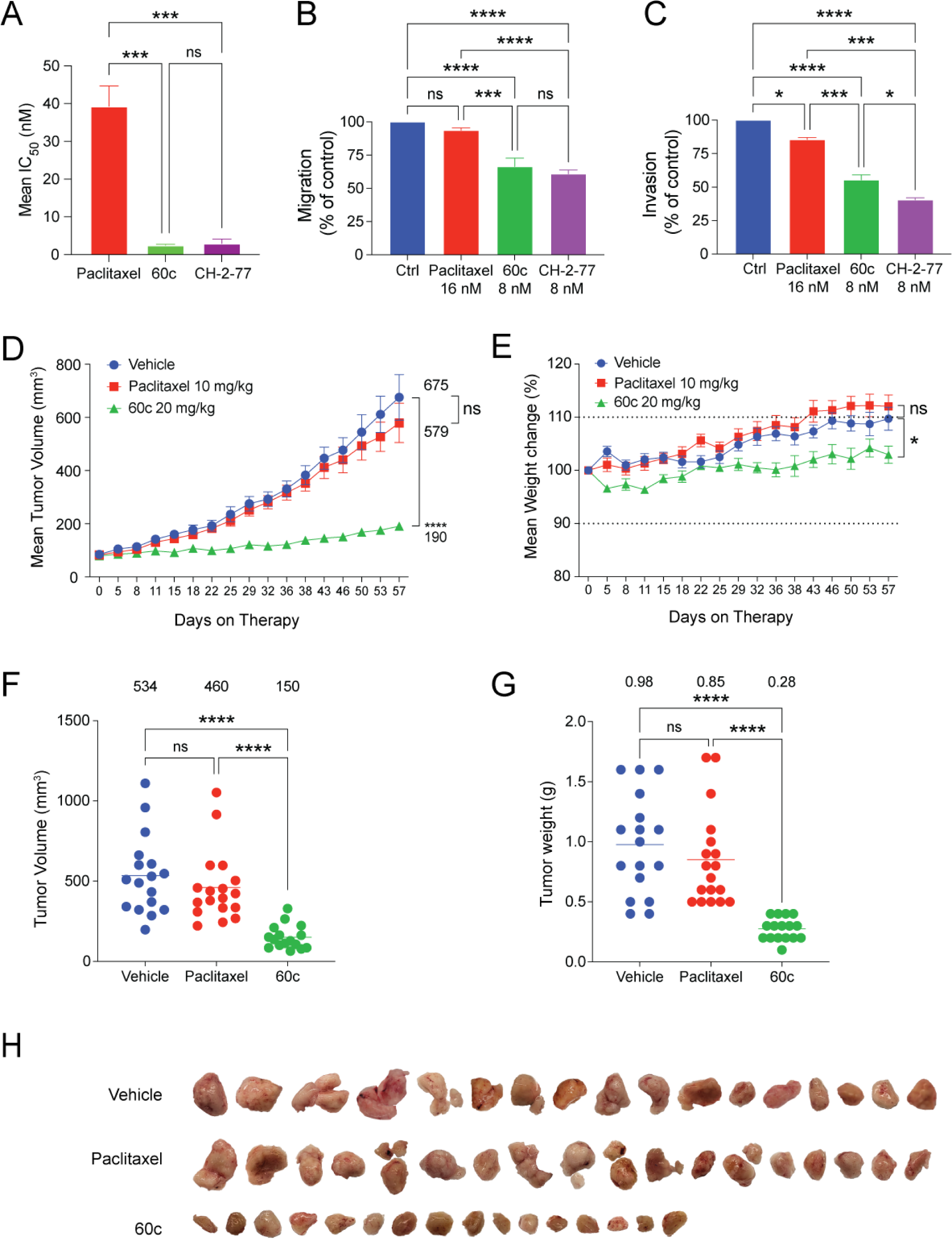
Efficacy of CH-2-77 and 60c versus paclitaxel in HCI-10-Luc2-PDX-derived cultured cells and head-to-head comparison of 60c to paclitaxel in HCI-10-Luc2 xenografts. (**A**) The IC_50_ values for paclitaxel, CH-2-77 and 60c were determined by the MTS assay in adherent PDX-cells derived from the HCI-10-Luc2 model. The bar graphs represent the grand mean ± SEM of the calculated IC_50_ values from three independent experiments, also refer to Table 2. The mean migration (**B**) and invasion (**C**) indices were determined after exposure to each compound for 24 h. Migration and invasion data are expressed as the grand mean of two independent biological experiments (n=3 technical replicates/experiment). (**D**) The mean tumor volume ± SEM over the course of therapy in response to vehicle, 10 mg/kg paclitaxel, IP (3x/week) or 60c (20 mg/kg, 5x/week, IP) treatment. Beginning on day 15 of therapy, the differences among all cohorts are significant by two-way ANOVA analysis. (**E**) The mean percent change in mouse body weight ± SEM during therapy relative to the starting body weight at day 0. Dashed lines indicate changes either 10% above or below the baseline (y-axis =100%). (**F**) Scatter plot of *ex vivo* final tumor volume, and final tumor wet weight (**G**). The mean ± SEM is indicated for each group. (**H**) Images of excised tumors. For all experiments, statistically significant differences were determined by one-way ANOVA followed by Dunnett’s multiple comparison tests at experimental endpoint; only pairwise analyses that are significant are shown; * *p* <0.05; ** *p* <0.01; **** *p* <0.0001; ns, not significant.

**Figure 7.**
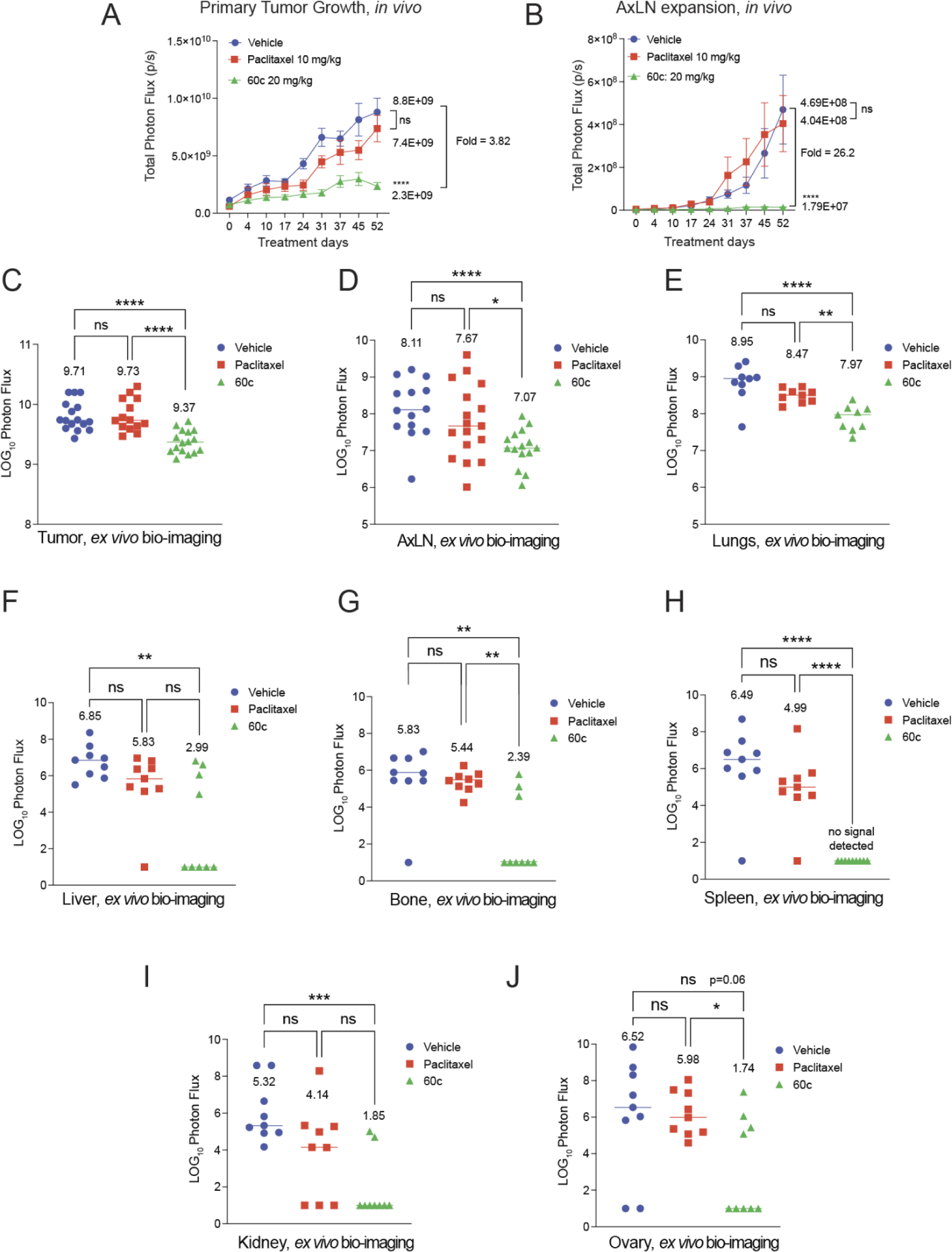
60c represses growth of pre-established axillary lymph node lesions and inhibits lungs, leg bones, spleen and kidney metastasis. (**A**) The mean total photon flux ± SEM was measured in primary tumors over the course of treatment, including all animals all shown in Figure 6. (**B**) Animals were covered with black paper so that light signal from the AxLN region could be imaged from intact mice; data are plotted as the mean total photon flux ± SEM. AxLN metastases were present at the time of randomization into treatment cohorts on day 0, allowing for measurement of the expansion of AxLN signal over therapy. The fold-change between day 0 and endpoint photon flux is indicated by brackets. (**C-J**) Excised primary tumors (**C**) and multiple organs (**D-J**) were immediately bio-imaged *ex vivo*, including the AxLNs (**D**), the lungs (**E**), the liver (**F**), the leg bones, cleaned of connective tissue (**G**), the spleen (**H**), the kidney (**I**) and the ovaries (**J**). For panels C-J, all data are plotted on a log_10_-transformed scale, wherein a value of 1.0 indicates no signal was detected. The mean ± SEM of total photon flux per organ is indicated above each group.

**Table 2.** IC_50_ (nM) ± SEM values for paclitaxel, 60c and CH-2-77 in HCI-10-Luc2 PDX-derived cells, calculated by the MTS assay. Data represent the grand mean of IC_50_ values derived from at least three biological experiments.

|  | **Paclitaxel** | **60c** | **CH-2-77** |
| --- | --- | --- | --- |
| HCI-10-Luc2 | 39.3 ± 5.4 | 4.5 ± 1.8 | 4.4 ± 1.1 |

A distinct advantage of the HCI-10-Luc2 PDX model is that tumor cells metastasize from the m.f.p. to the AxLNs before primary tumors are large enough to assign to treatment cohorts [16]. Therefore, the expansion of pre-established AxLN metastatic disease is measurable *in vivo* in intact mice throughout treatment. 60c significantly suppressed AxLN growth while on therapy, whereas AxLN metastases increased >20-fold in vehicle- and paclitaxel-treated mice (**Fig. 7B**). Overall, paclitaxel failed to inhibit primary tumor growth or metastasis to any tissue since there were no significant differences in total photon flux for any organ relative to the vehicle group (**Figs. 7C-7J**). In contrast, 60c significantly reduced metastatic burden in the AxLN, lungs, liver, leg bones, spleen, kidney, and ovary (**Figs. 7D-7J**). Notably, no animals in the 60c cohort developed detectable metastases in the spleen by bio-imaging (**Fig. 7H**). Furthermore, several individual mice within the 60c treatment cohort did not develop detectable metastases in the liver (5 of 9; 55.6%), bone (6 of 9; 67%), kidney (7 of 9; 78%), and ovary (5 of 9; 56%) (also refer to **Supp. Fig. 7** for organ-specific frequencies). When compared with previously collected data using VERU-111 treatment in the same HCI-10-Luc2 PDX model [16], we noted a pattern of metastatic tissue tropism among the CBSIs. First, VERU-111 strongly represses liver metastases; no detectable photon flux was observed in livers of VERU-111-treated mice [16]. In contrast, 44% of mice receiving 60c developed liver metastasis; the differences in mean photon flux between vehicle- and 60-treated mice were only significant if livers without signal were included in the 60c group (**Supp. Fig. 7A**). 60c also inhibited leg bone metastases more effectively than VERU-111. Only 3 of 9 mice had detectable signal in either set of leg bones after 60c treatment, while all mice treated with VERU-111 showed detectable bone metastasis signal (**Fig. 7G** and **Supp. Fig. 7B**). Furthermore, whereas VERU-111 had minimal effect on spleen or kidney metastasis, no mice receiving 60c exhibited splenic metastasis signal (**Fig. 7E and Supp. Fig. 7C-D**). As another measure of the number of animals per cohort impacted by metastasis (presence or absence), a Fisher’s exact test was used for each organ site comparing 60c to VERU-111. This analysis revealed a significant difference in the frequency of mice impacted by metastasis for the liver, spleen, kidney, and leg bones for 60c (**Table 3**). Conversely, VERU-111 treatment was only significantly correlated with metastatic repression at the liver relative to 60c (**Table 3**). Next, a drug distribution study was performed using equivalent doses of 60c (IP) and VERU-111 (oral, route used previously to treat the HCI-10-Luc2 PDX model). Overall, 60c had good drug penetration in all organs, but was qualitatively superior to VERU-111 for the spleen, kidney, and brain (**Supp. Fig. 7F-G**). 60c was also retained in blood plasma at ∼9-fold higher concentration than VERU-111.

**Table 3:** Fisher’s exact tests were used to compare frequencies of observed organ-specific metastasis in the HCI-10-Luc2 PDX model for 60c- and on VERU-111-treated mice. Data are summarized from the current study with 60c and from previously reported VERU-111 data (https://doi.org/10.1158/1535-7163.MCT-19-0536). Significant *p*-values are bolded.

| 60c treated mice |  |  |  |  | VERU-111 treated mice |  |  |  |
| --- | --- | --- | --- | --- | --- | --- | --- | --- |
|  | Mets | Vehicle | 60c | *p*-value | Mets | Vehicle | VERU-111 | *p*-value |
| Liver | Negative | 0 | 5 | **0.0294** | Negative | 0 | 6 | **0.0022** |
|  | Positive | 9 | 4 |  | Positive | 6 | 0 |  |
| Spleen | Negative | 1 | 9 | **0.0004** | Negative | 8 | 10 | 0.6728 |
|  | Positive | 8 | 0 |  | Positive | 4 | 3 |  |
| Kidney | Negative | 0 | 7 | **0.0023** | Negative | 5 | 9 | 0.2377 |
|  | Positive | 9 | 2 |  | Positive | 7 | 4 |  |
| Bone | Negative | 1 | 6 | **0.0498** | Negative | 0 | 0 | >0.9999 |
|  | Positive | 8 | 3 |  | Positive | 10 | 11 |  |

## 4. Discussion

Taxanes remain a frontline agent for metastatic breast cancer patients, but rapid onset of chemoresistance and common adverse events remain major roadblocks in patient quality of life. Developing novel effective agents against taxane-refractory tumors, including CBSIs like VERU-111 and its derivatives, is a crucial next step. Potency in the sub-nM range is desirable for CBSIs to be potential payloads in ADCs for targeted TNBC therapy [44].

Modifications were made to the VERU-111 structure to improve potency and metabolic stability. 60c and CH-2-77 each showed enhanced binding to the tubulin macrostructure and improved cell growth inhibition over VERU-111 in a panel of conventional TNBC cell lines. Like other tubulin inhibitors, CH-2-77 and 60c induced expression of the apoptotic markers cleaved PARP, cleaved caspase-3, and phosphorylated BCL2, 60c induced apoptosis at a concentration as low as 8 nM.

CH-2-77 and 60c were each more potent than paclitaxel to prevent wound closure in parental TNBC cells, and were also efficacious in TxR models at a concentration as low as 8 nM. In transwell migration assays using 231-TxR cells, both drugs were highly potent at 10 nM, a significant improvement over VERU-111, which was ineffective in TxR cells up to 15 nM. In addition, 10 nM of either 60c and CH-2-77 inhibited 231-TxR cell invasion through extracellular matrix.

In the 231-TxR xenograft model, CA-4P, 60c and CH-2-77 efficiently suppressed primary tumor growth and lung metastasis, with no discernable differences among the three CBSIs. In contrast, tumor growth in the paclitaxel cohort overlapped with the vehicle control group. Paclitaxel partially suppressed lung metastasis, although not as effeciently as the CBSIs. It is possible that paclitaxel is partially effective to repress 231-TxR lung metastasis because circulating tumor cells have lost cell survival signals from the extracellular matrix (anoikis) and from neighboring cells. As expected, 60c induced apoptosis and reduced cell proliferation; however, none of the CBSIs changed microvessel density.

In the more aggressive, taxane-refractory HCI-10-Luc2 PDX model, 60c not only reduced PDX tumor growth and tumor photon flux, but also prevented the expansion of AxLN metastatic disease. In contrast, paclitaxel had no significant effect, even with increased frequency of paclitaxel administration (3x/week, IP @ 10 mg/kg) used in the current study vs. the previous study (1x/week, IV @ 10 mg/kg) [16].

All PDX-bearing mice develop lung metastasis, and 60c was equipotent to VERU-111 to inhibit lung metastases [16]. However, 60c appeared to exhibit distinct anti-metastatic tissue tropism compared to VERU-111, which was previously evaluated in the same HCI-10-Luc2 PDX model. For example, 60c potently suppressed bone metastasis, with only 3 of 9 mice exhibiting bone signal, whereas all mice treated with VERU-111 developed some degree of bone metastasis [16]. In contrast, whereas VERU-111 completely inhibits liver metastasis [16], half of the mice receiving 60c therapy developed liver metastasis, suggesting that VERU-111 outperforms 60c in the context of liver metastasis.

The HCI-10-Luc2 PDX model also facilitates evaluation of CBSI efficacy on the more rare splenic and renal metastatic niches that are typically observed in late-stage patients [45–49]. In this study, most mice receiving vehicle (8 of 9) or paclitaxel (8 of 9) developed splenic metastases, yet no bioluminescent signal was detected in the spleens of 60c-treated mice (0 of 9 mice). We cannot compare splenic efficacy between the 60c data herein and the previous VERU-111 results because in the current study, ∼90% of vehicle-treated mice developed splenic metastases, whereas fewer than 25% of mice developed splenic lesions in the prior study [16]. Most mice bearing HCI-10-Luc2 tumors also develop kidney metastasis. Only 2 of 9 mice had kidney signal with 60c treatment whereas only 4 of 13 mice had kidney signal with VERU-111, indicating likely similar efficacy. Drug distribution analyses showed that more 60c than VERU-111 was present in the spleen and kidney, suggesting either improved tissue penetration or reduced efflux at these sites could be partly responsible for the enhanced efficiency of 60c. We also observed enrichment of VERU-111 relative to 60c in the kidneys and liver, possibly explaining VERU-111’s efficacy against liver metastases. These studies are qualitative, and the mechanism and specificity of tissue penetration and blood plasma retention of 60c relative to other CBSIs warrants further investigation.

Clinicians have limited options for stage IV TNBC, particularly for taxane-refractory disease [50]. Although there have been recent advances in therapies for TNBC patients with immune checkpoint inhibitors in the neoadjuvant setting (Keytruda, NCT03036488) [51], and therapies that target germline BRCA mutations, few breast cancer patients are candidates for these therapies. It will likely be necessary to continue to incorporate broad-spectrum anti-proliferative agents like tubulin inihibitors in treatment regimens. We have shown that 60c potently blocks cell proliferation and cell motility/metastatic spread in both taxane-sensitive and taxane-resistant TNBC lines and in two metastatic, TxR xenograft models. We have also observed an apparent tissue tropism in metastatic efficacy between 60c and VERU-111, which requires further investigation into each compound’s PK/PD profiles in tumor-bearing animals to determine if bio-distrubution or tissue-specific drug metabolism explains these results. Unlike VERU-111, 60c was detected in brain tissue, so new derivatives of 60c could be rationally designed to facilitate penetration of the blood-brain barrier, making it an attractive treatment option for patients with brain metastases. Novel derivatives of 60c could be generated to further improve potency to sub-nM IC_50_ values to facilitate development of novel antibody-drug conjugates using 60c as the warhead. Overall, our data position 60c as a promising candidate for future clinical trials for patients with late-stage TNBC unresponsive to the current standard of care, and for patients with multi-site extracranial metastatic burden.

## Author contributions

**Conception and design:** D. Oluwalana, K.L. Adeleye, R. I. Krutilina, T.N. Seagroves, W. Li

**Development of methodology:** T.N. Seagroves, W. Li

**Acquisition of data (provided animals, acquired and managed patients, provided facilities, etc.):** D. Oluwalana, K.L. Adeleye, R.I. Krutilina, H. Chen, H. Playa, S. Deng, D.N. Parke, J. Abernathy, L. Middleton, A. Cullom, D.N. Parke, B.Thalluri, D. Ma, H.C. Playa, T.N. Seagroves, W. Li

**Analysis and interpretation of data (e.g., statistical analysis, biostatistics, computational analysis):** D. Oluwalana, K.L. Adeleye, R.I. Krutilina, H. Chen, H. Playa, S. Deng, D.N. Parke, B.Thalluri, D. Ma, B. Meibohm, T.N. Seagroves, W. Li

**Writing, review, and/or revision of the manuscript:** D. Oluwalana, K.L. Adeleye, R.I. Krutilina, D.D. Miller, T.N. Seagroves, W. Li

**Administrative, technical, or material support (i.e., reporting or organizing data, constructing databases):** D.D. Miller, T.N. Seagroves, W. Li

**Study supervision:** T.N. Seagroves, W. Li

## Acknowledgements

Live-cell imaging was supported through the UTHSC Center for Cancer Research molecular imaging shared resource. We thank Dr. Alana Welm of the Huntsman Cancer Institute for providing the HCI-10 PDX. The research reported in this publication utilized resources obtained from the Preclinical Research Shared Resource at Huntsman Cancer Institute at the University of Utah, supported by the National Cancer Institute of the National Institutes of Health under Award Number P30CA042014.

## Funding

This work was supported by the National Institutes of Health (CA148706 to W.L. and D.D.M.), the Department of Defense Breast Cancer Research Program (W81XWH2010011 to W.L. and W81XWH2010019 to T.N.S), the University of Tennessee Drug Discovery Center, and the UTHSC Office of Research new grant support award. The contents of the article are solely the responsibility of the authors and do not necessarily represent the official views of the NIH/NCI or the DoD.

## Notes

### Competing Interest Statement

W.L. and D.D.M. previously received sponsored research agreements (SRAs) from Veru, Inc., although the SRAs are not directly related to this study. W.L. is scientific consultant for Veru, Inc. However, Veru, Inc. did not have any influence on this study, and only required reviewing this study prior to submission for publication for potential intellectual property purposes.

