## Supplementary Figure 1 for "Biological activity of a stable 6-aryl-2-benzoyl-pyridine colchicine-binding site inhibitor, 60c, in metastatic, triple-negative breast cancer"

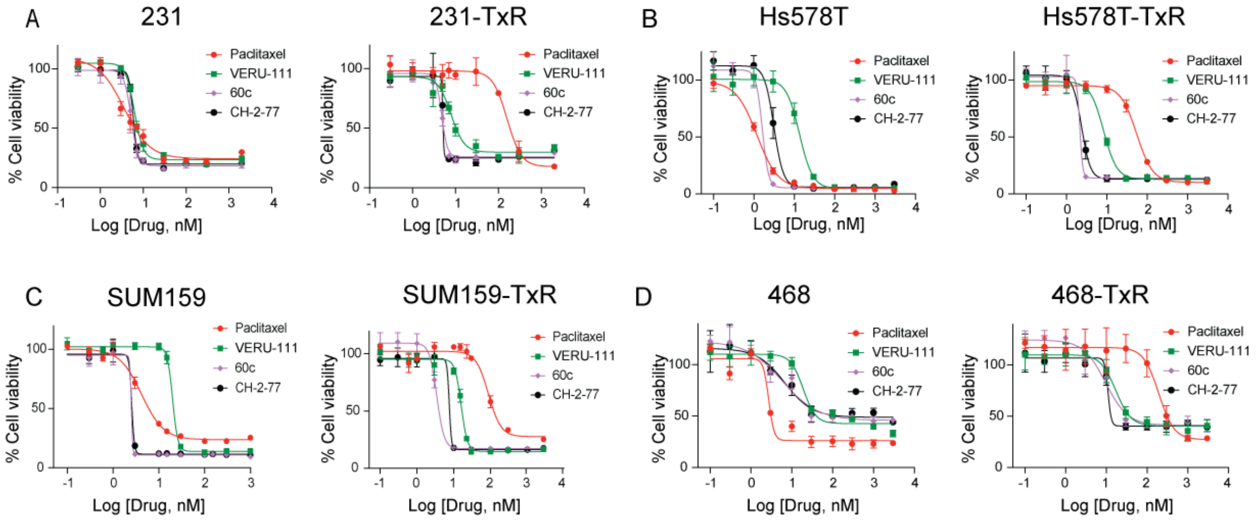


**Supplementary Figure 1. Growth inhibitory effect of 60c and CH-2-77 measured by the MTS assay**. The growth inhibition effect of 60c and of CH-2-77 relative to paclitaxel and VERU-111 was determined after 48 h (231, Hs578T, SUM159, SUM159-TxR), 72h (231-TxR, Hs578T-TxR) or 96h (468, 468-TxR) of drug incubation over a range of concentrations (0.3 nM to 3.0 μM) and expressed as cell viability (%) at study endpoint. Representative response curves of three independent biological replicates are shown for parental and taxane-resistant 231 (**A**), Hs578T (**B**), SUM159 (**C**), 468 (**D**) cells.
