## Supplementary material for "Biological activity of a stable 6-aryl-2-benzoyl-pyridine colchicine-binding site inhibitor, 60c, in metastatic, triple-negative breast cancer": Supplmentary Figure 2

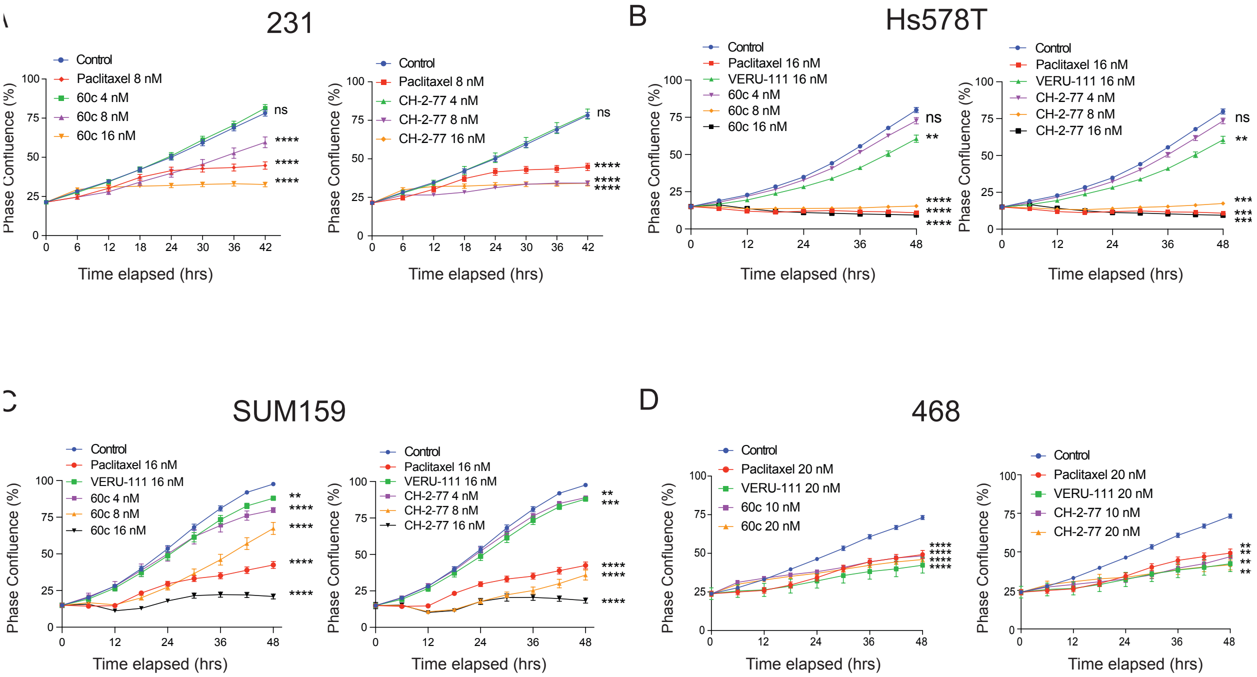


**Supplementary Figure 2. 60c and CH-2-77 significantly suppress parental TNBC cell growth in a dose dependent manner, measured in real-time by cell confluence.** Parental 231 (**A**), Hs578T (**B**), SUM159 (**C**) and 468 (**D**) cells were exposed to paclitaxel, VERU-111, 60c or CH-2-77 for 42 h (231), 48 h (SUM159, Hs578T or 468). Drugs were added at the indicated final concentration, and the medium was not changed over the course of the experiment. Dose-dependent changes in growth were measured by live cell imaging using the IncuCyte S3 imager and quantified by the percent (%) confluence algorithm. A representative drug response curve for at least three independent biological replicates per cell line is shown; *** *p* <0.001; **** *p* <0.0001; ns, not significant.
