## Supplementary Figure 3 for "Biological activity of a stable 6-aryl-2-benzoyl-pyridine colchicine-binding site inhibitor, 60c, in metastatic, triple-negative breast cancer"

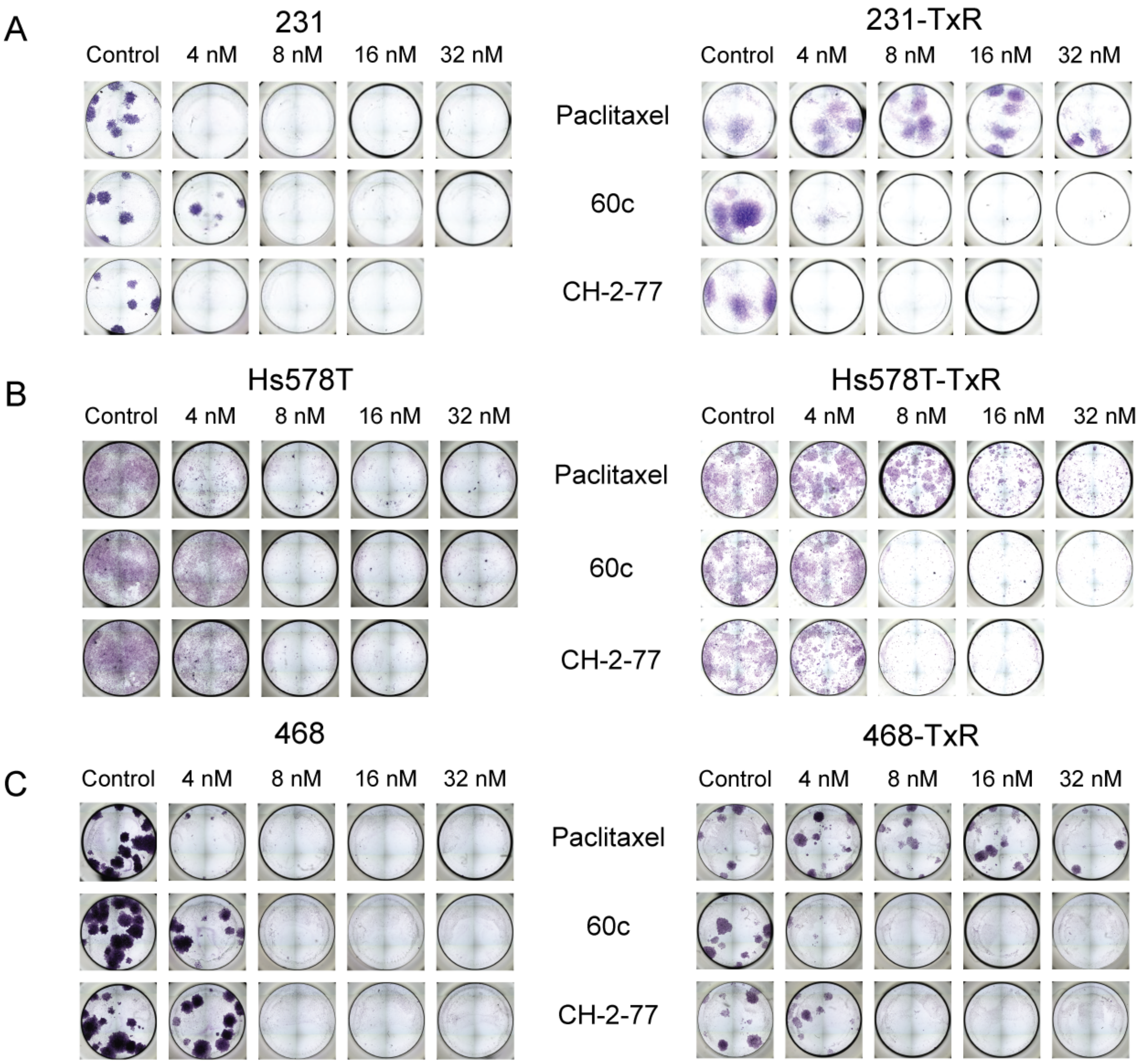


**Supplementary Figure 3. 60c is comparable to CH-2-77 to inhibit colony growth in taxane-sensitive and taxane-resistant TNBC cells.** Images of colony formation assays at endpoint when parental or TxR sublines of 231 (**A**), Hs578T (**B**), or 468 (**C**) cells are treated with increasing concentrations of paclitaxel, 60c or CH-2-77 (4 - 32 nM). All well images are representative of technical replicates (n = 8, vehicle or n = 4, each compound). Paclitaxel was only effective against parental TNBC cells, inhibiting clonogenicity at 4 nM in those lines, but it had minimal effect on the TxR sublines.
