## Supplementary Figure 4 for "Biological activity of a stable 6-aryl-2-benzoyl-pyridine colchicine-binding site inhibitor, 60c, in metastatic, triple-negative breast cancer"

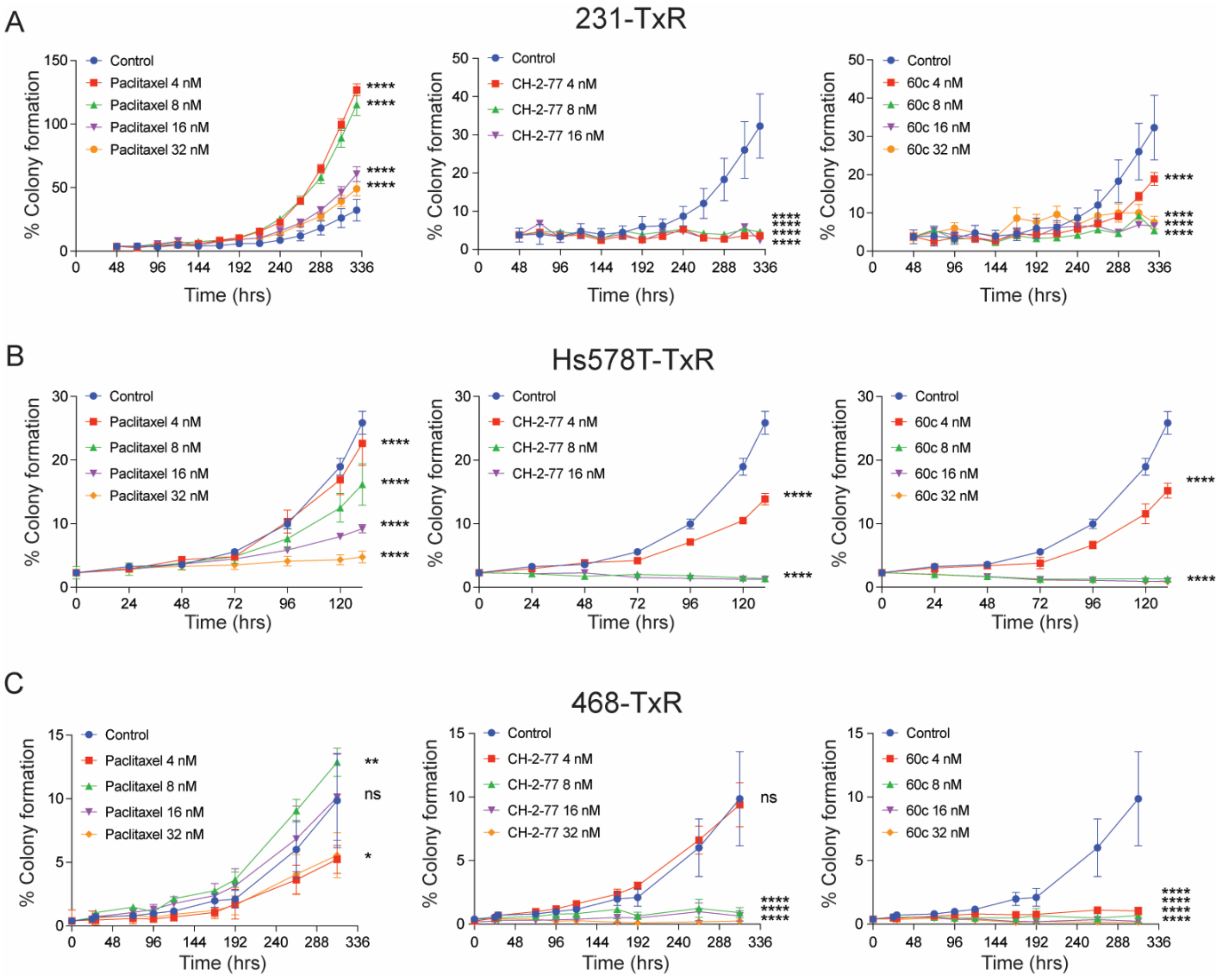


**Supplementary Figure 4. 60c and CH-2-77 suppress colony formation of TNBC TxR sublines at low nanomolar concentrations.** Clonogenic growth of 231-TxR (**A**), Hs578T-TxR (**B**) and 468-TxR (**C**) cells was measured for at least 5 days in the presence of either paclitaxel, 60c or CH-2-77 in a range from 4 nM to 32 nM. Assay duration was based on the increased doubling time of TxR cells relative to parental cells. The area occupied by cells was determined in real-time using the phase density algorithm of the IncuCyte S3 imaging system. Data were transformed to reflect the percentage (%) of colony formation relative to time 0 h (or 48 h for 231-TxR). Data representative of two biological replicate assays are shown. The mean ± SEM of % colony formation is shown over time (n = 8 wells, vehicle; n = 4 wells for each drug). Data were analyzed by two-way ANOVA followed by Dunnett's multiple comparisons testing. All p-values reflect the comparison to the vehicle control at experimental endpoint; * $p$ <0.05; ** $p$ <0.01; **** $p$ <0.0001; ns, not significant.
