## Supplementary Figure 5 for "Biological activity of a stable 6-aryl-2-benzoyl-pyridine colchicine-binding site inhibitor, 60c, in metastatic, triple-negative breast cancer"

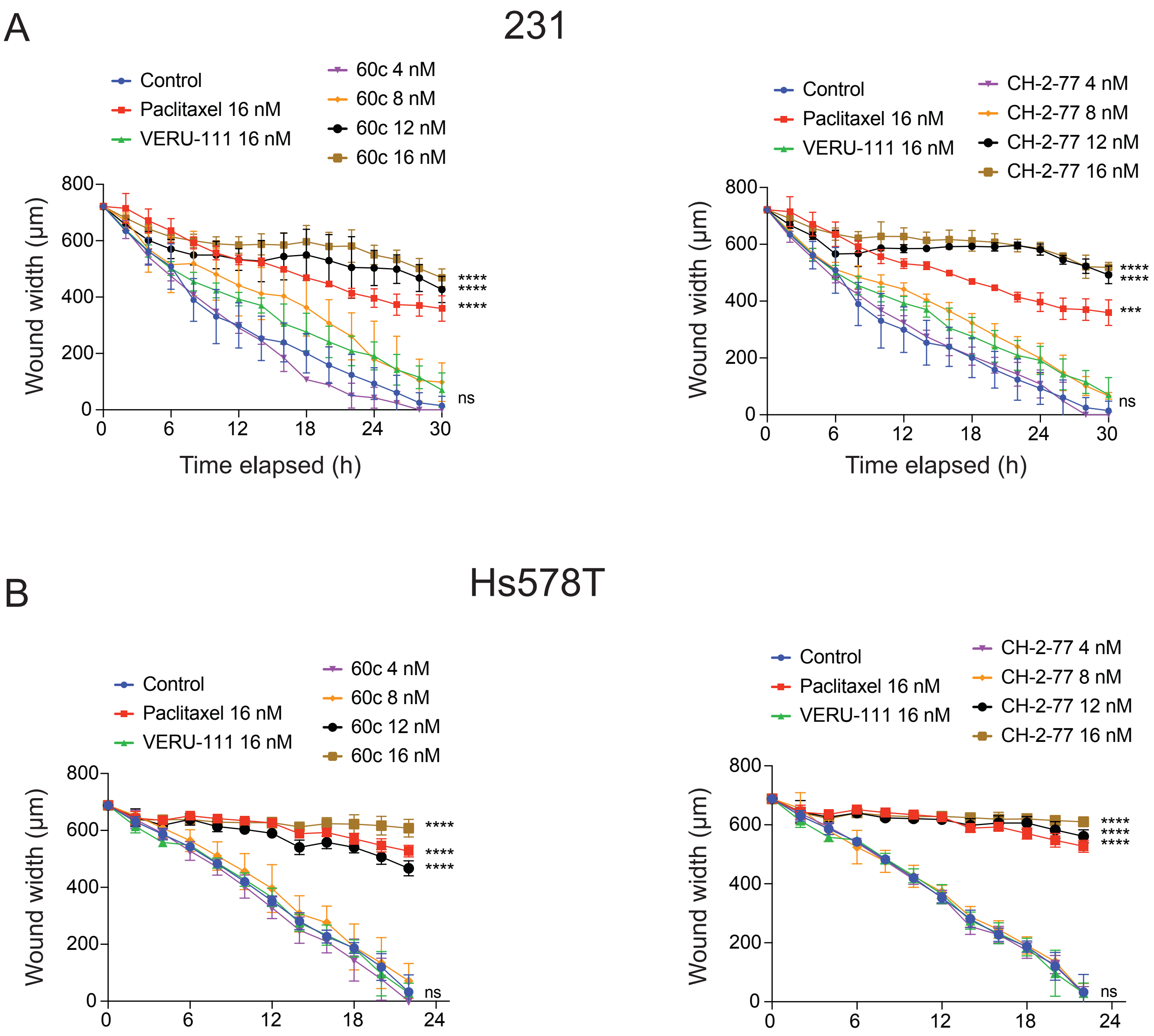


**Supplementary Figure 5. Dose-dependent inhibition of wound healing by 60c and CH-2-77 in parental, taxane-sensitive TNBC cells.** The rate of wound closure in the presence of paclitaxel (16 nM), VERU-111 (16 nM), or increasing concentrations of 60c and CH-2-77 (4, 8, 12 or 16 nM) was measured in 231 (**A**) and Hs578T (**B**) parental cells using the IncuCyte S3 live-cell imager as in Figure 3. Data were analyzed by two-way ANOVA followed by Dunnett's multiple comparisons testing. All *p*-values are shown relative to the vehicle control group at the experimental endpoint; ** $p < 0.01$; *** $p < 0.001$; **** $p < 0.0001$; ns, not significant.
