## Supplementary Figure 6 for "Biological activity of a stable 6-aryl-2-benzoyl-pyridine colchicine-binding site inhibitor, 60c, in metastatic, triple-negative breast cancer"

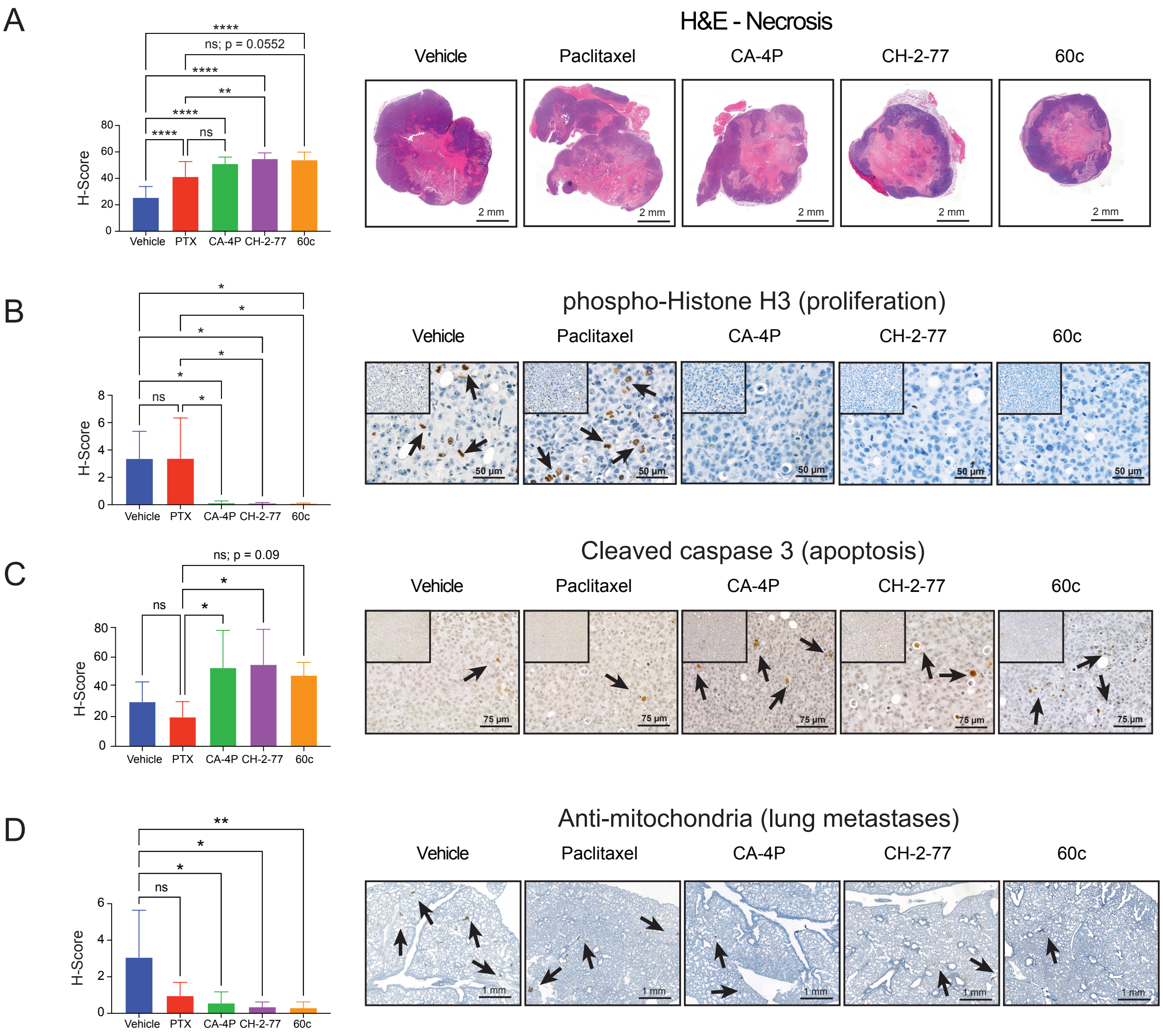


**Supplementary Figure 6. Histological analysis of orthotopic 231-TxR tumors confirms that CBSIs induce tumor cell death, while suppressing proliferation and metastatic spread.** (**A-D**) Whole tumor sections were stained with H&E (**A**) or immunostained with (**B**) phospho-histone H3(Ser10) (RRID:AB_331535), or (**C**) cleaved caspase-3 (RRID:AB_234118) and signal intensity quantified using the H-score algorithm in the 3D-Histech QuantCenter per the materials and methods. Lung metastasis burden was evaluated after immunostaining sections with a human specific mitochondrial marker (RRID:AB_10562769) (**D**). All data were analyzed by one-way ANOVA and post-hoc Tukey's multiple comparison testing; * *p* <0.05; ** *p* <0.01; **** *p* <0.0001; ns, not significant.
