## Supplementary Figure 7 for "Biological activity of a stable 6-aryl-2-benzoyl-pyridine colchicine-binding site inhibitor, 60c, in metastatic, triple-negative breast cancer"

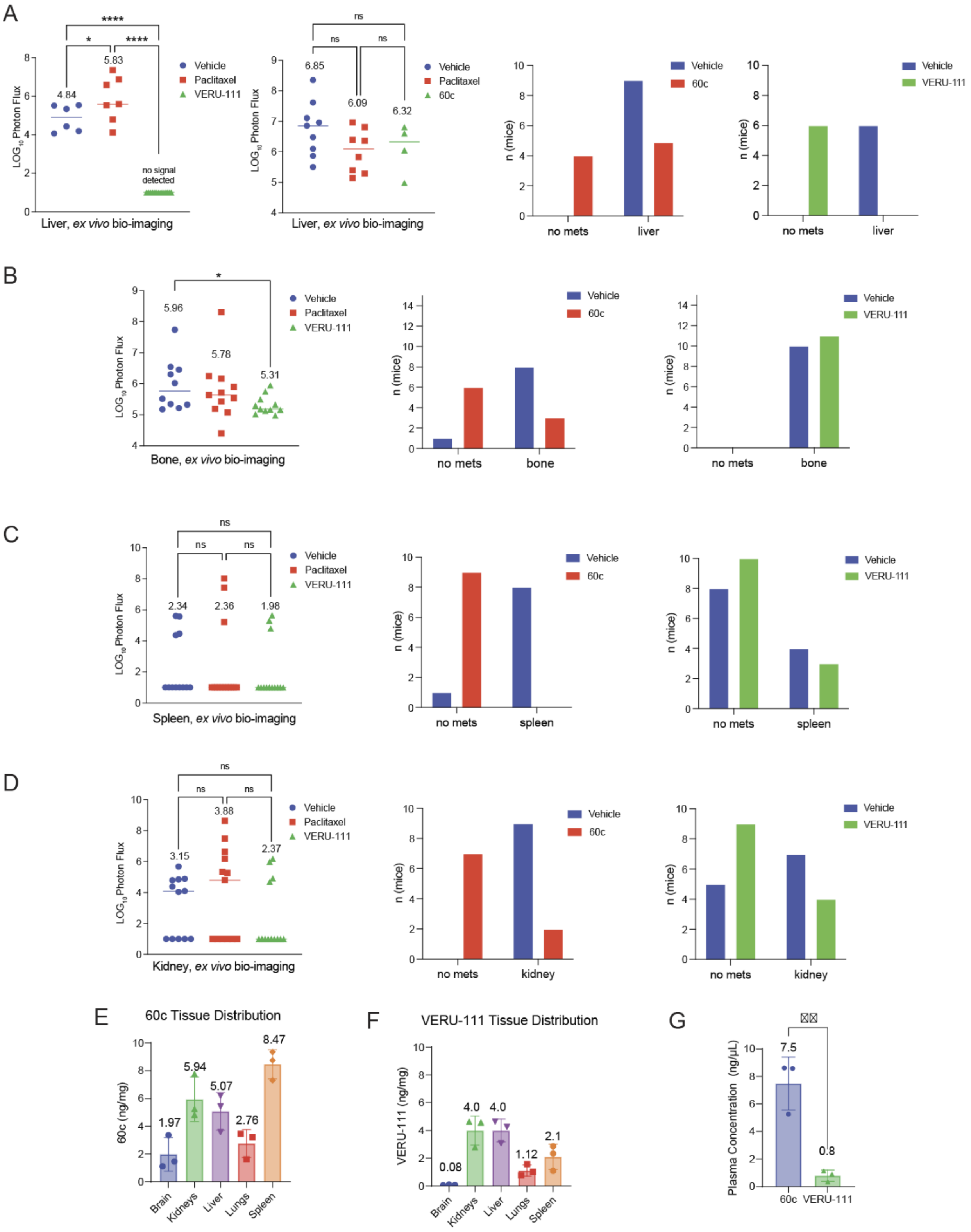

A
B
C
D
E
F
G
Liver, ex vivo bio-imaging
Bone, ex vivo bio-imaging
Spleen, ex vivo bio-imaging
Kidney, ex vivo bio-imaging
Vehicle
Paclitaxel
VERU-111
60c
no signal detected
no mets
liver
bone
spleen
kidney
n (mice)
LOG10 Photon Flux
60c Tissue Distribution
VERU-111 Tissue Distribution
60c (ng/mg)
VERU-111 (ng/mg)
Plasma Concentration (ng/μL)
Brain
Kidneys
Liver
Lungs
Spleen

**Supplementary Figure 7. Comparison of *ex vivo* bio-imaging of organs and drug distribution in the HCI-10-Luc2 PDX model after treatment with VERU-111 or 60c**. Scatter plot points represent indvidual *ex vivo* imaged organs and the mean is indicated above each column. All data are expressed on a $\log_{10}$ scale, unaffected tissues with no detectable light signal have a value of 1.0. Bar graphs represent the number mice per group observed with detectable metastases for indicated organs after 60c or VERU-111 treatment (**A**) As we previously reported (doi.org/10.1158/1535-7163.MCT-19-0536), VERU-111 significantly suppressed liver metastasis in all treated animals. In contrast, only 4 mice (50%) treated with 60c did not develop liver metastasis. Analyzing signal intensity in liver for all signal-positive mice, there was no significant difference in photo flux between vehicle, paclitaxel and 60c-treated animals. (**B**) As we previously reported, VERU-111 significantly represses metastatic expansion in the leg bones, but has no significant effect on spleen (**C**) or kidney (**D**) metastasis. Panels A and C were reproduced from [25]. Tissue distribution of 60c (IP, **E**) and VERU-111 (PO, **F**) 30 minutes or 1 h after administration, respectively, of a 25 mg/kg dose, and the blood plasma concentration of 60c and VERU-111 (**G**) at the time of tissue harvest**.**
